# A Chromosome Scale Genome Assembly for the Australian blue mussel *Mytilus planulatus*

**DOI:** 10.64898/2026.10.28.762448

**Authors:** Dona Kireta, Daniela Eugenia Nerelli, Amaro Saco, Enrico Bortoletto, Samantha Howitt, Umberto Rosani, Cynthia Riginos, Marco Gerdol

## Abstract

Mussels belonging to the genus *Mytilus* are among the world’s most ecologically and economically important bivalve molluscs. Although the genus most likely originated in the Northern Hemisphere, the evolutionary history of Southern Hemisphere lineages, including the timing and number of colonisation events leading to the extant Australian populations, remains debated. Limited genomic resources have hindered the reconstruction of these evolutionary histories. In Australia, distinguishing native *Mytilus planulatus* from invasive *Mytilus galloprovincialis* is critical for biosecurity and conservation, but previous molecular approaches lacked sufficient resolution to resolve species boundaries and hybridisation patterns.

Here, we present the first chromosome-scale reference genome for *M. planulatus*. The assembly consists of 14 chromosomes and over 51,000 annotated protein-coding genes per haplotype. Comparative analysis revealed extensive structural variation and widespread hemizygosity despite a largely conserved chromosomal organisation relative to *M. galloprovincialis*. Population analyses using SNPs and gene presence-absence variation in resequenced individuals confirmed previously inferred patterns of population structure and establish this reference genome as a valuable resource for future studies of genome evolution, adaptation and population genomics in *Mytilus*.

## DATA DESCRIPTION

### Background information

*Mytilus* mussels are ecologicallyand economicallyimportant ecosystem engineers, widely distributed in temperate coastal marine environments of both the Northern and Southern Hemispheres. They are farmed across their global range, represented a traditional food source in many coastal regions since ancient times, and play key ecological roles as reef-building, filter-feeding organisms that contribute to water purification [1, 2, 3].

From an ecological perspective, *Mytilus galloprovincialis* mussels are among the most successful marine invaders, and anthropogenic transport through aquaculture and shipping has facilitated repeated introductions of non-native species both in regions where no native *Mytilus* species were present (e.g. South Africa) and in regions where native mussel populations were already locally established (e.g. Chile, Eastern Asia and the Pacific coast of North America) [4, 5].

There are seven distinct *Mytilus* species that are often interfertile (sometimes referred to as the *Mytilus edulis* species complex) [6]. Despite extensive efforts, the taxonomy and evolutionary history of these species remain partially unresolved. In the Northern Hemisphere, different species within the *M. edulis* complex occasionally display distribution overlaps, creating large natural hybrid zones, resulting in widespread introgression [7, 8]. Similar scenarios are, however, also possible in the Southern Hemisphere, where native and invasive populations recently introduced from the Northern Hemisphere coexist. The difficulty in recognising easily detectable species-specific shell morphology traits, together with the scarcity of genetic resources for native Southern Hemisphere species, has long represented a challenge for species identification in these areas.

In Australia, species identification is particularly problematic as both the native mussel *M. planulatus*, and *M. planulatus × M. galloprovincialis* hybrids can be found, prompting biosecurity concerns [9, 10]. Current evidence supports *M. planulatus* as the native species, with pure populations found in southern Australia and Tasmania, while *M. galloprovincialis*, a globally invasive species whose native range spans the Mediterranean Sea and the Atlantic coasts of Western Europe, is thought to have established hybrid populations in regions such as Sydney Harbour and southern New South Wales (NSW) (Howitt in preparation) [9, 10]. Importantly, Atlantic and Mediterranean lineages of *M. galloprovincialis* were more divergent from each other than either was to *M. planulatus*, and the ancestry of the admixed *M. galloprovincialis × M. planulatus* differed between locations [9]. Until recently, these considerations have largely been based on the analysis of a limited number of molecular systems, such as the analysis of mitochondrial markers [11], allozymes [12], and a small number of polymorphic nuclear loci [13, 10]. In the past few years, however, the use of -omic approaches, such as RNA-sequencing, has provided an important confirmation to previous observations [9]. While informative, these approaches lack the resolution needed to fully disentangle hybridisation patterns and species boundaries in such a complex context. A high-quality reference genome for *M. planulatus* therefore represents a critical resource for supporting future studies in validating species identity, resolving the evolutionary relationships between this native species and its Northern Hemisphere relatives, and investigating invasion dynamics and adaptation in Southern Hemisphere *Mytilus* populations.

### Sampling, DNA extraction and sequencing

Wild mussels were sampled from a native *M. planulatus* population in the bay of Port Arthur, Tasmania (latitude: -43.14723, longitude: 147.85283) [10, 9]. Tasmania was chosen because previous population genomic analyses found Tasmanian *M. planulatus* to represent the endemic lineage, with no evidence of admixture with *M. galloprovincialis* [9]. Six adult individuals of undetermined sex were sampled from a jetty, and kept fresh until dissection. Sampled tissues (i.e. mantle, gill and posterior adductor muscle) were subsequently preserved at -80 °C until further processing. One of the sampled individuals was selected for the generation of the genome assembly. The collected gill tissue from this individual was split into two equal parts. The sample was in parallel subjected to high molecular weight DNA extraction, for the subsequent preparation of a PacBio HiFi library (following the manufacturer’s instructions), and for the construction of a Hi-C library using the Arima-HiC kit and standard workflows. The mantle tissue of the five remaining samples collected in Port Arthur, together with five individuals each from two additional populations from Rushcutter’s Bay in Sydney, NSW, and Eden Wharf in southern NSW, underwent whole genome resequencing (WGR). The Sydney mussels were expected to show admixed ancestry with Mediterranean *M. galloprovincialis*, whereas mussels from southern NSW were expected to show admixed ancestry with Atlantic *M. galloprovincialis*, based on previous population genetic studies [9]. These latter 10 samples were first preserved in ETOH, then transported on dry ice to the sequencing facility.

The PacBio HiFi library was sequenced on a PacBio Revio platform, while the Hi-C library and the 15 WGR libraries were sequenced on a DNBSEQ-T7 platform, using 150 bp paired-end sequencing. Raw PacBio reads were processed by the sequencing facility using the standard PacBio HiFi circular consensus sequencing workflow to generate high-fidelity consensus reads. The resulting HiFi reads were passed through quality control and filtering steps, which included adapter contamination, and low-quality read removal. After filtering, 560,794,526 Hi-C read pairs were retained, comprising 168.24 Gb of sequence data, with 97.04% of bases having a Phred quality score greater than Q20. A total of 4,010,202 HiFi reads were retained after filtering, comprising 78 Gb of sequencing data, corresponding to approximately 56*×* coverage, with a mean read quality of 32.5, a mean read length of 19,450 bases, and a read length N50 of 19,770 bases. Library preparation and sequencing were carried out by BGI Genomics (Hong Kong).

### Genome assembly and evaluation

PacBio HiFi reads were first assembled *de novo* using Hifiasm (v0.25.0-r726)[14] generating two partially phased pseudohaplo-type assemblies, hereafter referred to as haplotype 1 and 2 for simplicity. Both were evaluated using QUAST (v.5.3.0)[15], which indicated that both haplotypes had comparable assembly statistics and quality metrics. Assembly completeness was estimated using BUSCO (v.6.0.0)[16] against the metazoa_odb12 lineage dataset and using a k-mer based approach through Merqury (v.1.3)[17].

Both haplotype assemblies were then scaffolded with the Hi-C reads using YaHS (v.1.2.2)[18], after which assembly quality was re-evaluated with QUAST, BUSCO and Merqury. Hi-C contact maps were generated using PretextMap (v.0.1.9)[19] and visualised with PretextSnapshot (v.0.0.5)[20].

The initial assembly consisted of 1.42 Gb with 717 contigs and an N50 of 6.2 Mb for haplotype 1, while haplotype 2 comprised 1.41 Gb with 538 contigs and an N50 of 6.6 Mb. Both haplotypes demonstrated high completeness and very low duplication, fragmentation, and missingness rates, indicated by BUSCO scores of 99.3% (single-copy = 97.5%, duplicated = 1.8%, fragmented = 0.2%, missing = 3) for the first and 99.0% (single-copy = 98.2%, duplicated = 0.7%, fragmented = 0.2%, missing = 5) for the second (Fig. 1). Merqury confirmed the near-complete representation of the genome in the final assembly, since the two haplotypes showed k-mer completeness values of 69.37% and 68.86%, respectively, corresponding to an overall completeness of 99.37%, while the median quality value (QV) reached 67.68 (Fig. 2). K-mers exclusively associated with the reads but absent from the assembly were restricted to low multiplicity values, consistent with sequencing errors rather than unassembled genomic sequence, and represented only a minor fraction of the total k-mer space. Haplotype-specific k-mers peaked at the expected haploid coverage depth, indicating correct assignment of haplotype specific sequences to their respective assemblies, while k-mers shared between haplotypes peaked at approximately twice the haploid coverage, as expected for sequences shared by both haplotypes. The well-defined and clearly separated peaks observed in the k-mer distribution graphs further supported the high completeness and low error rate of both assembled haplotypes.

**Figure 1.**
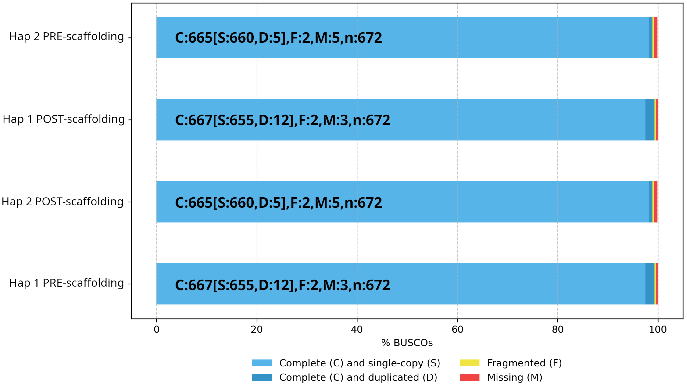
BUSCO completeness scores estimated using the metazoa_odb12 lineage dataset before and after scaffolding.

**Figure 2.**
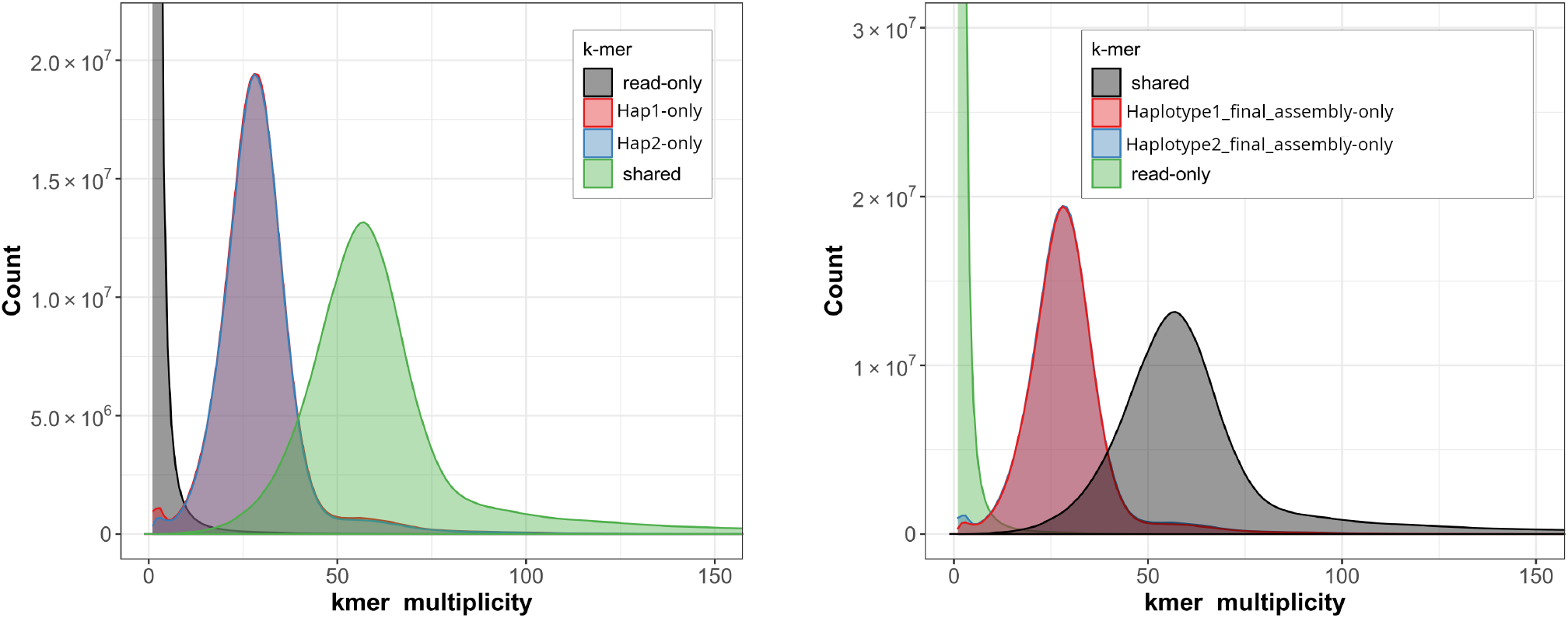
Merqury spectra of haplotype (hap) 1 and 2 assemblies before scaffolding (left) and after scaffolding (right) illustrating the k-mer multiplicity distribution across both haplotype assemblies. In the left panel grey represents k-mers found exclusively in the reads, red represents k-mers exclusive to the first haplotype, blue represents k-mers exclusive to the second and green represents shared k-mers. After scaffolding, in the right panel, green represents k-mers found exclusively in the reads, red and blue represent k-mers exclusive to haplotype 1 and two respectively, while grey represents k-mers shared between the two haplotypes.

After Hi-C scaffolding with YaHS, assembly contiguity improved, increasing the N50 to 99.9 Mb for haplotype 1 and 105.3 Mb for haplotype 2 (Fig. 3). Additional contiguity metrics further validated the quality of the scaffolded assembly, with an N90 of 77.6 Mb (L90: 13) for the first and 74.81 Mb (L90: 13) for the second haplotype. The final assembly of haplotype 1 spanned 1.4 Gb, with 97.86% of the sequence assigned to 14 chromosome-level scaffolds. Haplotype 2 was characterised by similar metrics, with 98.45% of the sequence associated with the 14 largest pseudochromosomal scaffolds. Hi-C contact maps (Fig. 3) showed strong interaction signals along the diagonal with defined inter-chromosomal boundaries, confirming the organisation of the assemblies into 14 discrete chromosome-scale scaffolds and supporting their structural integrity. This is consistent with other *Mytilus* species, based on both karyotype characterisation and genome sequencing approaches [21, 22, 23, 24, 25, 26]. A comprehensive summary of all the assembly quality and completeness statistics is provided in Table 1.

**Table 1.** Quality evaluation of assembly assessed by Quast and Merqury.

| Assembly | Number of contigs | N50 (Mb) | N90 (Mb) | L50 | L90 | k-mer completeness (%) | Quality value (QV) |
|---|---|---|---|---|---|---|---|
| Haplotype 1 pre-scaffolding | 717 | 6.2 | 1.6 | 70 | 236 | 69.37 % | 66.86 |
| Haplotype 2 pre-scaffolding | 538 | 6.6 | 1.7 | 66 | 223 | 68.86% | 68.70 |
| Haplotype 1 | 364 | 99.9 | 77.6 | 7 | 13 | 69.37 % | 66.86 |
| Haplotype 2 | 196 | 105.3 | 74.8 | 7 | 13 | 68.86% | 68.70 |

**Figure 3.**
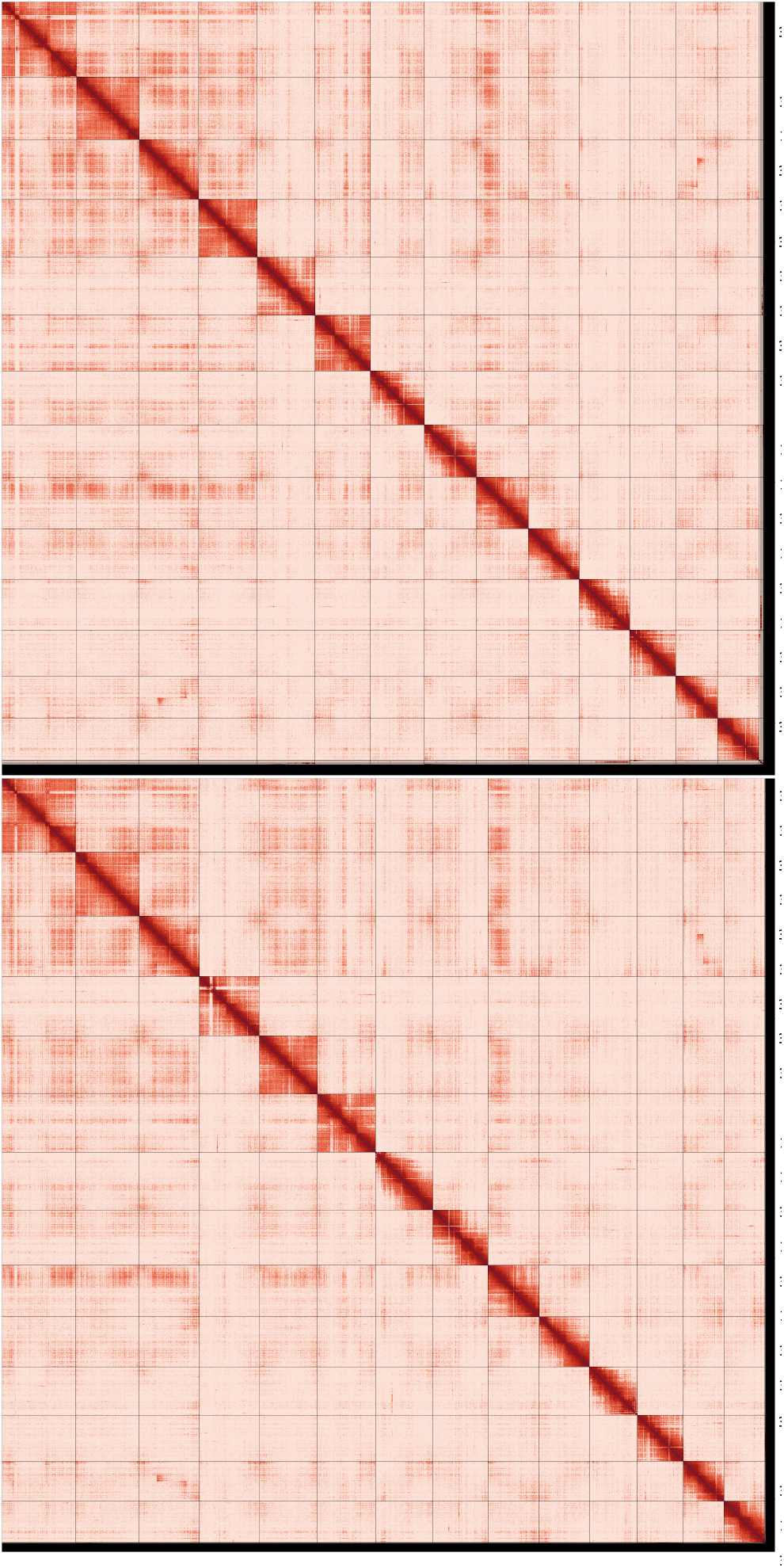
Hi-C contact maps of the primary (left) and alternate (right) assemblies that display the chromosomal interaction patterns across the 14 predicted pseu-dochromosomes. No major inversions, translocations, or large gaps are apparent, and the off-diagonal interaction blocks are consistent with expected chromatin folding patterns.

Finally, the assembly was screened for potential contamination using Flagger [27] and BlobToolkit [28]. HMM-Flagger identifies local depth anomalies indicative of collapsed, duplicated, or erroneous regions. The analysis classified 0.67% of the assembly as erroneous and 0.24% as duplicated. Potential contamination of the 14 chromosomes was additionally screened using BlobToolKit, revealing no contamination signal. Taken together, these results support the high quality of the genome assembly.

#### Gene annotation

To annotate repetitive regions in the *M. planulatus* genome, RepeatModeler (v.2.0.5)[29] was used with default parameters to generate a *de novo* repeat library. This was subsequently filtered to remove sequences shorter than 100 bp, and sequences in which unresolved nucleotides accounted for more than 5% of the repeat length. This novel collection of repeats was combined with Repbase repeat sequences and clustered using uclust to generate the final non-redundant repeat library, which was used as input for RepeatMasker (v.4.1.9)[30] to soft-mask repeats in the genome.

Gene annotation was then performed using BRAKER3 (v.3.0.8) [31, 32, 33, 34, 35, 36, 37, 38, 39, 40, 41, 42, 43, 44], a semi-unsupervised pipeline that integrates both RNA-seq expression data and protein homology evidence to guide gene prediction. RNA-seq evidence was derived from publicly available datasets of multiple *M. galloprovincialis* tissues, including gills, hemolymph, gonads, digestive glands and additional mixed tissue samples in order to capture genes with broader expression patterns, thereby improving annotation reliability (NCBI Sequence Read Archive accession numbers: SRR9984990, SRR9984953, SRR9984952, SRR9984951, SRR9984950, SRR1045900, SRR1046115, SRR11996735, SRR16641566, SRR16641571, SRR442032, SRR442036, SRR7044533, SRR7138611, SRR7138616, SRR7138628, SRR7138631, SRR7640807, SRR9984981 and ERR4296979). Protein homology evidence was based on the UniRef90 database [45] and on protein sets from closely related bivalve species including *Perna viridis, Modiolus philippinarum, Limnoperna fortunei, Magallana gigas, Pecten maximus, Mercenaria mercenaria, Tegillarca granosa* and *Bathymodiolus platifrons* in order to maximise the amount of available homology information and improved the detection of proteins. This combined approach predicted a total of 52,309 protein-coding genes for haplotype 1 and 51,449 protein-coding genes for haplotype 2. Annotation completeness was assessed using BUSCO against the mollusca_odb12 lineage dataset, with predicted protein set from haplotype 1 recovering 96.7% complete BUSCOs (94.5% single-copy, 2.2% duplicated), 0.4% fragmented, and 2.9% missing (Fig. 4).

**Figure 4.**
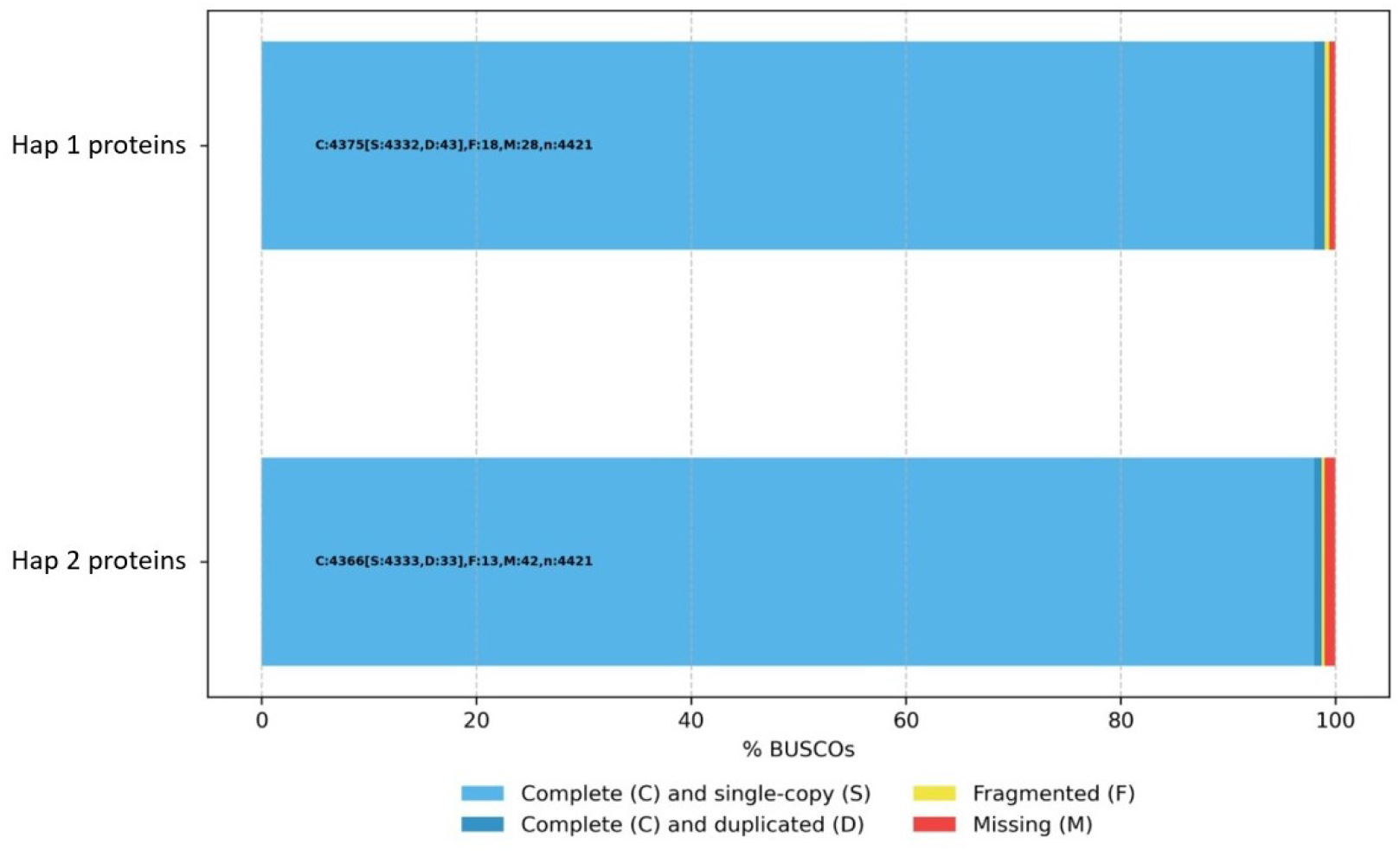
BUSCO completeness of the predicted protein sets for *M. planulatus* haplotypes 1 and 2, assessed using the mollusca_odb12 lineage dataset.

All software tools used in this study, along with their respective versions and repository links, are summarised in Table 2.

**Table 2.** Overview of bioinformatic tools, versions and repositories used for assembly.

| Software | Version | Source |
|---|---|---|
| PretextMap | 0.1.9 | https://github.com/sanger-tol/PretextMap |
| PretextSnapshot | 0.0.5 | https://github.com/wtsi-hpag/PretextSnapshot |
| BlobToolKit | 4.4.5 | http://github.com/genomehubs/blobtoolkit |
| RepeatModeler | 2.0.5 | https://github.com/Dfam-consortium/RepeatModeler |
| OrthoFinder | 3.1.5 | https://github.com/OrthoFinder/OrthoFinder |

#### Circos plot

To describe and visualize chromosomal characteristics of the *Mytilus planulatus* haplotype 1 assembly, a Circos plot was generated using Circos (v.0.69-8) [46]. The 14 chromosomes were divided into non-overlapping 1 Mb windows, and four metrics were calculated for each window: GC content, gene density, repetitive sequence density and mean gene dispensability. GC content was calculated as the percentage of G and C nucleotides in the window. Gene density was expressed as the number of annotated genes per Mb. Repeat density was calculated as the percentage of the window covered by annotated repetitive sequences. Gene dispensability was defined as the fraction of the 15 *Mytilus planulatus* individuals in which a gene was absent, based on the gene PAV matrix (see Gene PAV section below for details). The mean dispensability of all genes within each window was then calculated. These values were represented as separate tracks of the Circos plot, providing an overview of the distribution of genomic features and gene dispensability along the chromosomes (Fig. 5).

**Figure 5.**
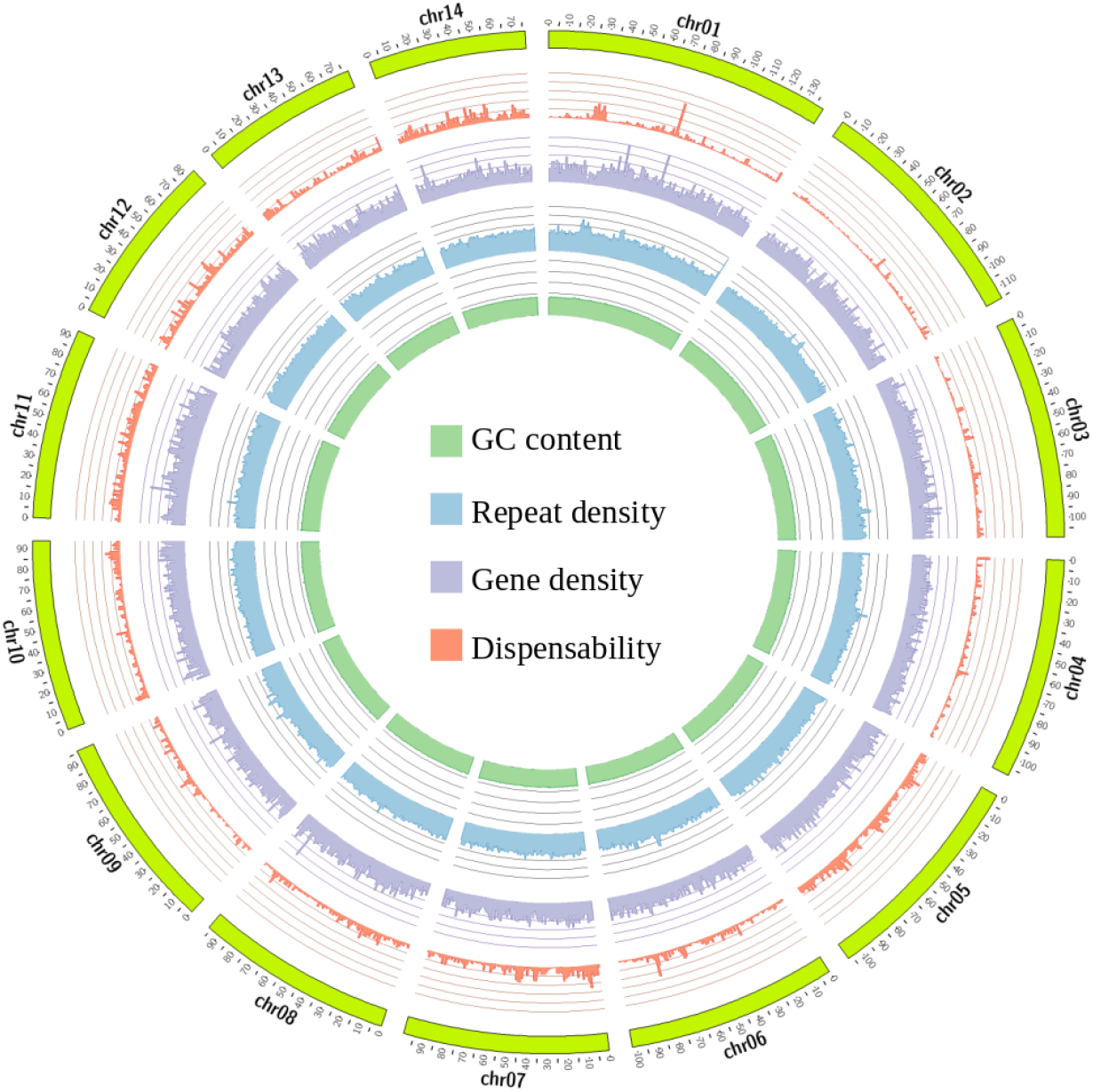
Circos plot of the *Mytilus planulatus* haplotype 1 assembly. From the outermost to the innermost track: the 14 chromosomes; mean gene dispensability per 1 Mb window (orange); gene density (purple), expressed as the number of annotated genes per megabase; repeat density (light blue), expressed as the percentage of each window covered by annotated repetitive sequences; and GC content (green), expressed as the percentage of G and C nucleotides in each window. The histogram tracks show the distribution of each metric along the chromosomes.

### Structural Variation analysis

To analyse structural variation (SV) in the *M. planulatus* genome, whole-genome alignments were produced between both *M. planulatus* haplotypes, and between the haplotype one of *M. planulatus* and the current NCBI reference haplotypes for *M. galloprovincialis* (GCF_965363235.1, sampling location Galicia, Spain) and *M. edulis* (GCF_963676685.1, sampling location Broad Haven, United Kingdom). These two references had comparable assemblies in terms of scale and contiguity, and were also comparable to our *M. planulatus* haplotypes. Both assemblies were chromosome-scale, comprising 14 chromosomes, and spanning 1.4 Gb of sequence. The *M. galloprovincialis* assembly had a scaffold N50 of 98 Mb, a contig N50 of 12.1 Mb, and 51*×* coverage. The *M. edulis* assembly had a scaffold N50 of 93.4 Mb, a contig N50 of 1.7 Mb, and 30*×* coverage.

For each comparison, chromosome correspondence was assessed using MashMap (v. 3.1.3), followed by whole genome alignments between homologous chromosomes with Minimap2 (v.2.28), using the asm20 mode, which allows for up to 10% divergence. The alignments were converted to BAM format and sorted using SAM-tools (v.1.21). Finally, SyRI (v.1.7) was used to define SV, and plotsr (v.1.1.1) for visualisation.

This analysis demonstrated a general conservation of structural synteny both between the two *M. planulatus* haplotypes, and between *M. planulatus* and the two reference species *M. edulis* and *M. galloprovincialis* (Fig. 6A), consistent with previous reports [47]. However, we also observed the presence of several large-scale rear-rangements, most notably intrachromosomal inversions. In addition, translocations and duplications were not negligible, although found at a lower frequency. Inversions were not uniformly distributed across the genome, with particularly high prevalence in some chromosomes, such as chromosome 11, which displayed significant structural rearrangements in all pairwise comparisons between haplotypes.

**Figure 6.**
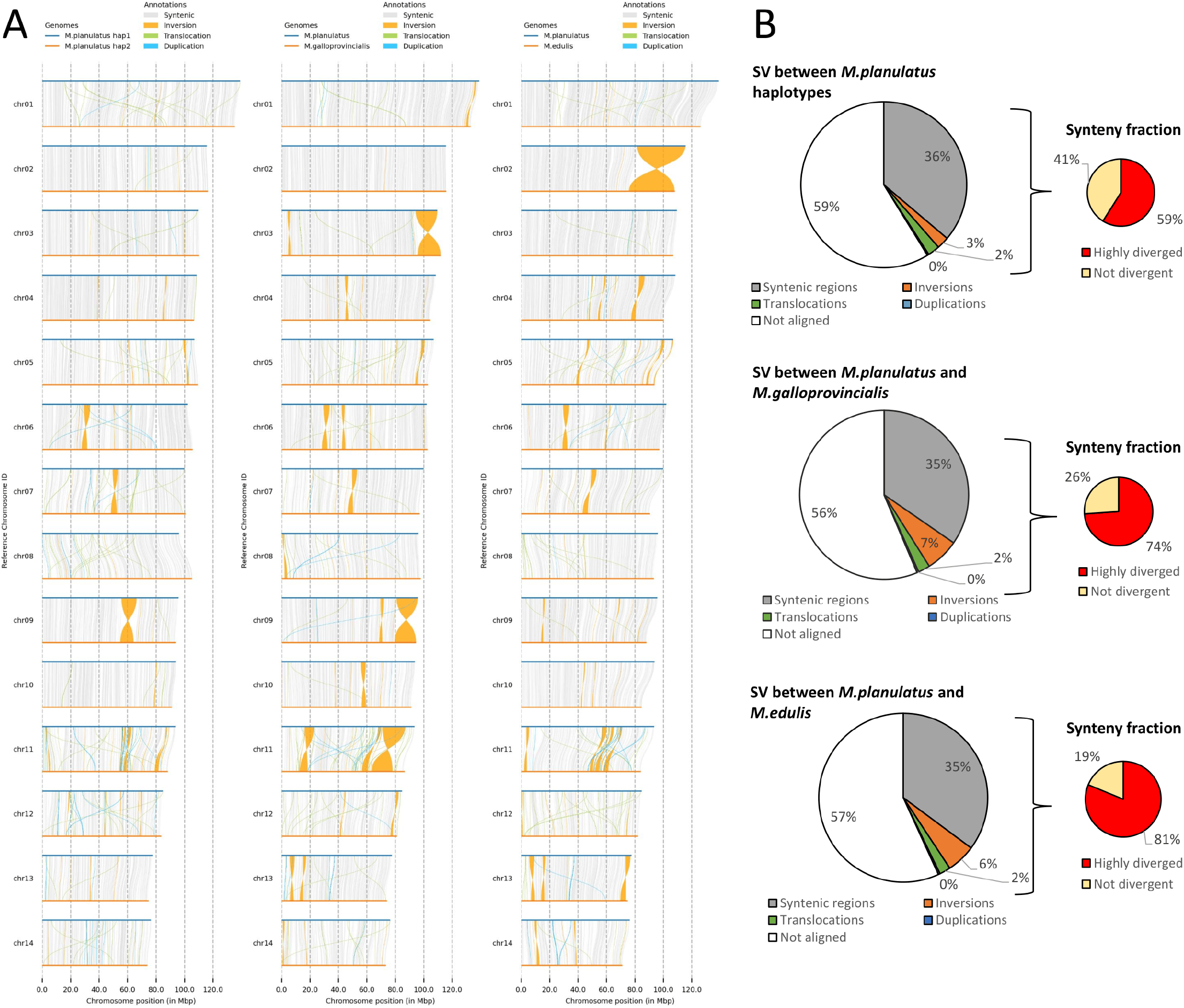
Structural variation between *M. planulatus* haplotypes, and *M. galloprovincialis* and *M. edulis* genomes. **A**: Whole genome alignments displaying structural variations; **B:** structural variation displayed as the fraction of total genome length for each of the pairwise genome alignments.

The most striking outcome of the SV analysis was that a significant fraction of the genome assembly, ranging between 56% and 59%, were not aligned between haplotypes (NOTAL sequences) (Fig. 6B). The fraction of misalignment did not change significantly in the comparison between the *M. planulatus* haplotypes, or between *M. planulatus* and two other mussel species, suggesting that these regions contain haplotype-unique sequences.

The high prevalence of haplotype-unique regions has been previously reported in other bivalves [47], and is believed to be due to the high hemizygosity reported in the genomes of *Mytilus* and other bivalve species, which is a hallmark of their pan-genomic nature [48, 49]. Although the unaligned fraction of the genome did not change between the structural comparisons, a smaller fraction of the aligned regions was classified as highly divergent between the two *M. planulatus* haplotypes (59%) than in the comparisons with *M. galloprovincialis* (74%) and with *M. edulis* (81%).

The NOTAL sequences identified between the two *M*.*planulatus* haplotypes were further analysed to estimate the real hemizygous fraction. After identifying gene synteny between the two haplotypes using MCScanX, an average sequence homology of 95% was found. This corresponded to the 5% divergence threshold defined by the P90 of syntenic alignments between the homozygous regions of the two haplotypes where these syntenic genes were located. Therefore, a 5% divergence threshold was established for confident cross-haplotype alignments. NOTAL sequences were then mapped to the alternate haplotype, and alignments exceeding the divergence threshold or covering less than 20% of the NOTAL sequence length were excluded. Read coverage analysis further supported this filtering, revealing that 44% of the NOTAL sequences were confidently haplotype-specific, while 38% were putative divergent sequences present in both haplotypes (Appendix 1 Figure 1).

After identifying gene synteny between the two haplotypes using MCScanX, a significant association (odds ratio=1.88; p-value=1.1 *×* 10^-266^) was found between structurally variable regions of the genome and haplotype-specific genes that lacked conserved synteny (accounting for 30% of all annotated genes), and between the syntenic aligned regions between the haplotypes and the syntenic genes (51% of all annotated genes). In addition, the homologies between these real allelic variants were consistently high (P85 of the alleles presented 95% homology or higher; P90 presented 90% homology or higher), which ensured avoiding important cross-mapping problems in the following PAV calculation, since the changes in mapping rate are insignificant at those divergence levels.

### Gene PAV and population context

#### Presence/Absence Variation (PAV)

The large SV found in the genome was further investigated by analysing the 15 resequenced *M. planulatus* individuals through a published gene PAV detection pipeline (https://github.com/Carmen-Tuc/PAV_pipeline), to categorise each gene either as core, if present in all individuals, or as dispensable, if absent in one or more individuals [50, 51]. In addition, population structure was investigated and compared to two previously published reference populations of *M. galloprovincialis* (likely the closest relative of *M. planulatus*) representative of the Atlantic and Mediterranean lineages [50], which consisted of 10 individuals from Ría de Vigo, Spain, and 6 individuals from the Goro Lagoon in Venice, Italy.

All reads from the 31 individuals (15 *M. planulatus* and 16 *M. galloprovincialis*) were mapped to haplotype 1 of the *M. planulatus* genome using `bwa-mem2` [52]. SAM files were converted to BAM format and indexed with SAMtools [53], and then processed with the PAV pipeline. Exon coordinates from the BRAKER annotation of haplotype 1 (see Gene Annotation section) were used to calculate normalised exon read coverage. These coverage estimates were then used to classify genes as present or absent in each individual relative to haplotype 1. For each sample, the expected coverage of single-copy genes was estimated from the BUSCO coverage distribution, and a gene was called present if its normalised coverage exceeded a threshold of 1/8 of the BUSCO coverage peak, and absent if coverage was below the threshold.

Across the 15 individuals mapped to that haplotype, 39,746 out of the 52,309 genes annotated in the *M. planulatus* haplotype 1 were present in ≥ 14 of the 15 samples (core genes), and 12,563 genes were categorised as dispensable (missing in ≥ 10% of individuals, i.e. ≥ 2). Note that although the definition of a core gene is that it occurs in every individual, we allowed a confidence margin to take into account the possibility of mapping errors. The PAV pipeline was also applied to the *M. galloprovincialis* reference individuals from the Atlantic and Mediterranean lineages to obtain comparable PAV patterns for the population-structure analysis. This allowed the construction of a binary PAV matrix spanning all 31 individuals.

#### Population structure

To explore and visualise population structure based on dispens-4 able gene content, the PAV matrix of all resequenced individuals was first filtered to retain only informative dispensable genes, defined as those present in more than 10% but fewer than 90% of individuals. The filtered matrix was then imported into R, where pairwise Jaccard distances between individuals were calculated using the `vegdist` function of the `vegan` package [54], and principal coordinate analysis (PCoA) was performed on the resulting distance matrix using the `pcoa` function from the ape package [55](Fig. 7A).

**Figure 7.**
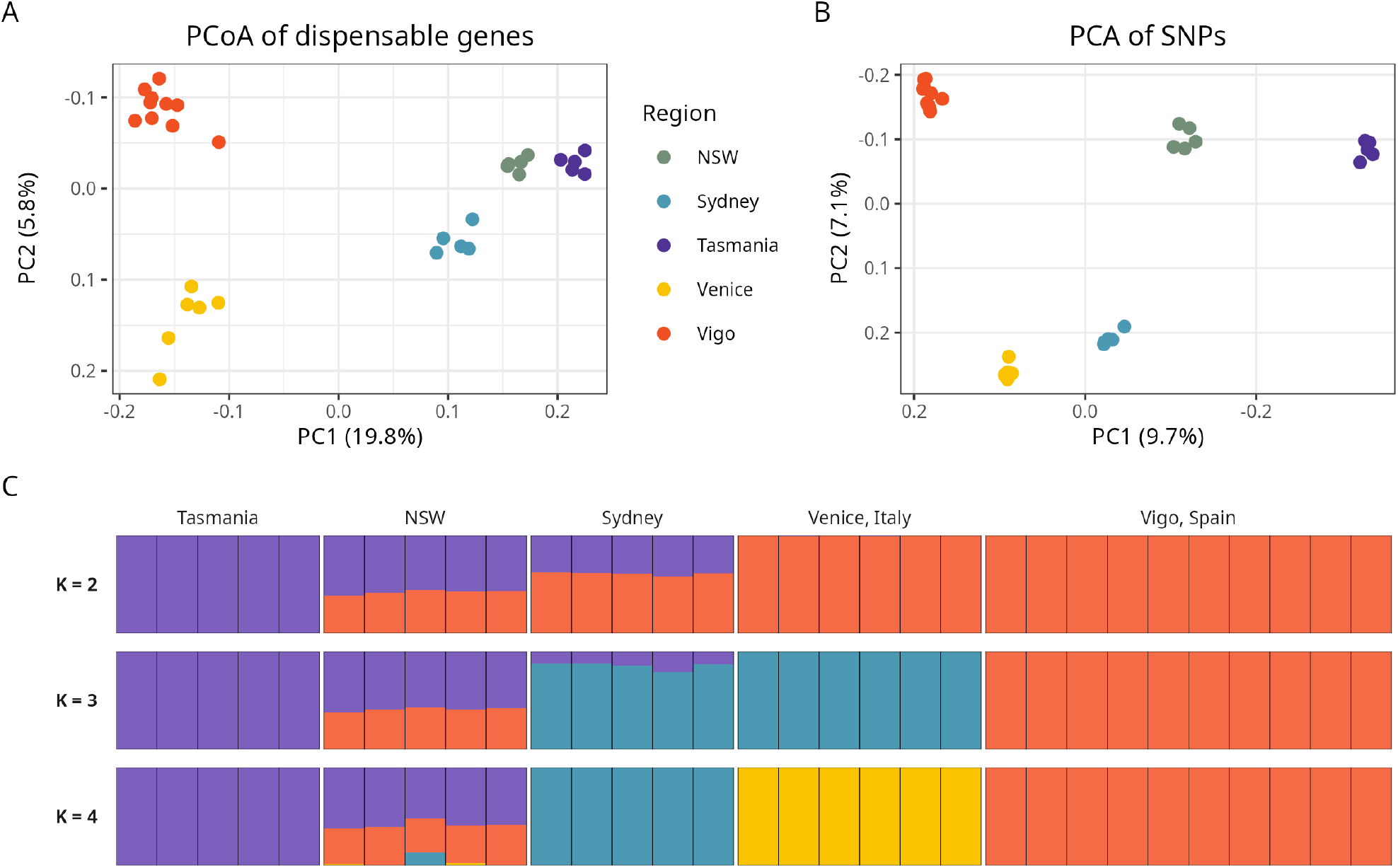
Population structure of *M. planulatus* from three Australian populations - Sydney, New South Wales, and Tasmania - versus two Northern Hemisphere reference populations of *M. galloprovincialis* from Venice in the Mediterranean Sea, and Vigo in the Atlantic Ocean. A: PCoA of dispensable genes based on Jaccard distance; B: PCA of SNPs; C: ADMIXTURE plots with k=2, 3 and 4.

To compare and corroborate the PAV population structure results with standard SNP-based variation, SNPs were also called and analysed. The mapped BAM files from all 31 individuals were duplicate-marked with Picard `MarkDuplicates` [56], and indexed with SAMtools. SNPs were called jointly using `bcftools mpileup, call` and `view`, by passing all samples into the same call, and providing population information with the -G flag [53], resulting in 203,749,316 SNPs. The variants were heavily filtered, to create a very high-quality final SNP set. Firstly, to remove SNPs that were poorly genotyped across the dataset, the merged VCF was split by population, and within each subset PLINK was used to retain only SNPs genotyped in at least 50% of individuals (`-geno 0.5`) as per the filtering strategy used by Nascimento et al. [57]. The union of the resulting per-population BED files was used as a regions file to filter the merged VCF with BCFtools `view -R`, retaining any SNP that passed the threshold in at least one population. Next, only biallelic SNPs found within core exons were kept, using a bed file of exon coordinates. Additional quality filters removed sites with low mapping and variant quality scores (`QUAL >= 30`), filtered using BCFtools `view`. Rare variants (minor allele frequency < 0.01) were filtered with PLINK. To reduce the effect of linked loci in the down-stream population genomic analysis, SNPs were further pruned for linkage disequilibrium in PLINK using a 50 kb sliding window with a one SNP step size, and r^2^ threshold of 0.1 (`–indep-pairwise 50kb` `1 0.1`). Of the initial 203,749,316 SNPs called, 6,646,004 remained after restriction to core exons and missingness filters were applied, and 115,084 SNPs were retained in the final MAF- and LD-pruned dataset used for PCA and ADMIXTURE analyses. This final dataset had an overall genotyping rate of 94%, with a mean per-individual genotype missingness of 5.6%. PCA was run using PLINK `–pca`, and ADMIXTURE analyses were run with K values of 2-4, additionally outputting the cross-validation errors. SNP-based PCA and admixture plots were visualised in R using `ggplot2` [58, 59] (Fig. 7B and 7C).

The SNP analysis was restricted to SNPs located within exons of core genes to mitigate biases arising from the highly hemizygous nature of *Mytilus* genomes. Since large portions of the genome are present in a hemizygous state, SNPs in these regions are expected to have high levels of missingness, which reflects SV rather than true allelic polymorphism, potentially confounding estimates of population structure. This approach ensured that only loci shared across all individuals were retained, allowing comparisons of homologous genomic regions, and reducing PAV-associated noise. Given the limited prior knowledge of *Mytilus* pangenome organisation and SV, such a strict variant filtering strategy was not possible in previous SNP-based population genetics studies carried out in *Mytilus*.

The population structure results based on the PAV pattern of informative dispensable genes (PCoA; Fig. 7A) and on SNPs (PCA; Fig. 7B) showed broad concordance, although the relative positions of the admixed Australian populations differed between analyses. The putatively pure *M. planulatus* population from Tasmania was most divergent from the *M. galloprovincialis* populations in both cases. In the SNP ordination, the resequenced individuals from the Sydney population, previously reported as strongly introgressed with the Mediterranean lineage of *M. galloprovincialis* [9], clustered very close to those belonging to the Mediterranean *M. galloprovincialis* population sampled in the Venice area. On the other hand, the southern NSW population, which is located geographically closer to Tasmania, occupied an intermediate position between the Tasmanian *M. planulatus* and Vigo *M. galloprovincialis* reference population, consistent with the previously hypothesised mixed ancestry with the Atlantic lineage of *M. galloprovincialis*. The PAV ordination largely supported the SNP structure, with the same broad differentiation, but showed a stronger separation between the Australian and European *M. galloprovincialis* populations. Along PC1, which explained approximately 20% of the variation, the Australian populations were positioned close together, and clearly separated from *M. galloprovincialis*. Tasmania and southern NSW were positioned particularly closely on this axis, while Sydney was located more intermediately. Along PC2, Sydney was positioned between the Australian and Mediterranean *M. galloprovincialis* populations, but was less strongly associated with the Mediterranean than in the SNP PCA.

The difference between the two analyses likely occurred because of a stronger reference bias in the PAV dataset. This analysis was re-4 stricted to the genes found in the *M. planulatus* haplotype 1 genome, therefore, by definition, all genes analysed were present in *M. planulatus*. Conversely, any *M. galloprovincialis* unique or PAV genes were not represented. In effect, the analysis only measured variation within the *M. planulatus* gene set. Any *M. galloprovincialis*-specific dispensable genes absent from the *M. planulatus* reference were excluded from the comparison, and we expect this increased the separation between *M. galloprovincialis* and *M. planulatus* in the PAV ordination. In contrast, the SNP dataset was heavily filtered and restricted to core exons conserved across all populations, so this form of reference bias had a lesser effect. These results therefore illustrate a limitation of PAV analyses for comparisons involving introgression and multiple species, and highlight the need for a comprehensive pangenome for future studies.

The ADMIXTURE analysis (Fig. 7C) supported and refined the ordination results. The cross-validation error was lowest at K = 2 (0.59), compared with K = 3 (0.65) and K = 4 (0.77), which indicated that K = 2 was the best supported model. At K = 2, two major ancestry components, corresponding to *M. planulatus* and *M. galloprovincialis*, were identified. Both the Sydney and NSW populations exhibited admixed ancestry, although *M. galloprovincialis* contributed the larger genomic component in Sydney (i.e. 61%), whereas *M. planulatus* predominated in the southern NSW population (i.e. 58%), consistent with the placement of these populations in both ordinations. At K = 3, the Atlantic (Vigo) and Mediterranean (Venice) *M. galloprovincialis* populations formed distinct ancestry components. The two admixed *M. planulatus* populations differed not only in the overall proportion of *M. galloprovincialis* ancestry, but also in its origin. Specifically, the Sydney population derived almost exclusively from the Mediterranean lineage (86% Mediterranean, 1% Atlantic, 14% *M. planulatus*), whereas the southern NSW population showed predominantly Atlantic *M. galloprovincialis* ancestry (41% Atlantic, 1% Mediterranean, 59% *M. planulatus*). Consequently, at both K = 2 and K = 3, the Sydney population consistently exhibited a greater overall contribution from *M. galloprovincialis* than the southern NSW population. These results are in close agreement with previously reported patterns of population structure across these regions [9, 10]. Finally, at K = 4, the Sydney population was assigned to a distinct ancestry component, whereas ancestry proportions in the southern NSW population remained largely unchanged. Across all K values, the putatively pure *M. planulatus* population from Tasmania consistently formed a distinct genetic cluster, indicating the great-5 est degree of differentiation from all other sampled populations.

### Functional enrichment of dispensable genes

To test whether the variable part of the genome was enriched for particular functions relative to the rest of the genome, the *M. planulatus* dispensable gene set (absent in more than 10% of individuals, i.e. at least 2 of 15), as defined by the PAV analysis, was tested against the full annotated gene set. Protein sequences were annotated with InterProScan [60] (v 5.63-95.0), to produce gene-to-PFAM mappings. Enrichment of each PFAM domain in the dispensable subsets was assessed using Fisher’s exact test [61]. Terms were considered significantly enriched at a Bonferroni-corrected p-value < 0.05. We further filtered terms associated with at least five annotated genes and at least three dispensable genes, and an observed-expected count difference > 4 or < -4, for significantly over- and under-represented annotations, respectively. Significantly enriched terms are reported in Appendix 1 Table 1.

Functional enrichment analysis of dispensable genes recovered many of the same gene families previously reported to be associated with PAV in *M. galloprovincialis* [48, 62]. This finding suggests that the functional repertoire affected by gene PAV is largely conserved between the two species. The recurrent enrichment of immune- and stress-related gene families is consistent with the hypothesis that these dispensable genes have been shaped by similar selective pressures during the evolutionary diversification of *Mytilus*, possibly reflecting adaptation to distinct environmental conditions and pathogen communities.

Some of the most strongly enriched domains in the set of dispensable genes were associated with transposable elements, which are among the main drivers of SV found in *Mytilus*, and others were involved in the immune response and adaptation. These include C1q-domain-containing proteins, fibrinogen-related proteins (FREPs), AIG1-domain GTPases (GIMAP/IMAP family), and MAB21-domain proteins. In contrast, C-type lectin, Toll/interleukin-1 receptor (TIR) and leucine-rich repeat (LRR) domains were significantly under-represented, with the latter two being typically associated with Toll-like receptors [48, 62]. Finally, several domains involved in the regulation of cell survival and growth were also significantly enriched, including caspase-activation and YAP domains, consistent with survival-related functions. A notable example was the enrichment of the AIG1-domain GTPases. This family has been previously linked to stress response and immune functions [63] and recently identified as over-represented in the dispensable genes of amphioxus [64].

### Correlation between structural variation and dispensable genes

The large hemizygous haplotype-unique sequences identified in the *M. planulatus* genome assembly (NOTAL) were filtered to identify the real hemizygous regions, as discussed in the Structural Variation Analysis section (Appendix 1 Figure 1). This hemizygous fraction is responsible for the pangenomic nature of the genome, since genes encoded in these regions are exclusively associated with one of the two assembled haplotypes, and therefore expected to be dispensable at the population level. To verify this, the location of the genes classified as dispensable by the PAV analysis was investigated in depth. The vast majority (95%) of dispensable genes were placed outside of the syntenic alignments, in genomic regions associated with SV between haplotypes. As seen in Figure 8A, dispensable genes were enriched in the general NOTAL fraction (1.9 odds ratio) but even more so in the putative hemizygous sequences identified (2.88 odds ratio). Meanwhile, the opposite occurred in the NOTAL regions that were found to be present but divergent in both haplotypes (0.39 odds ratio).

**Figure 8.**
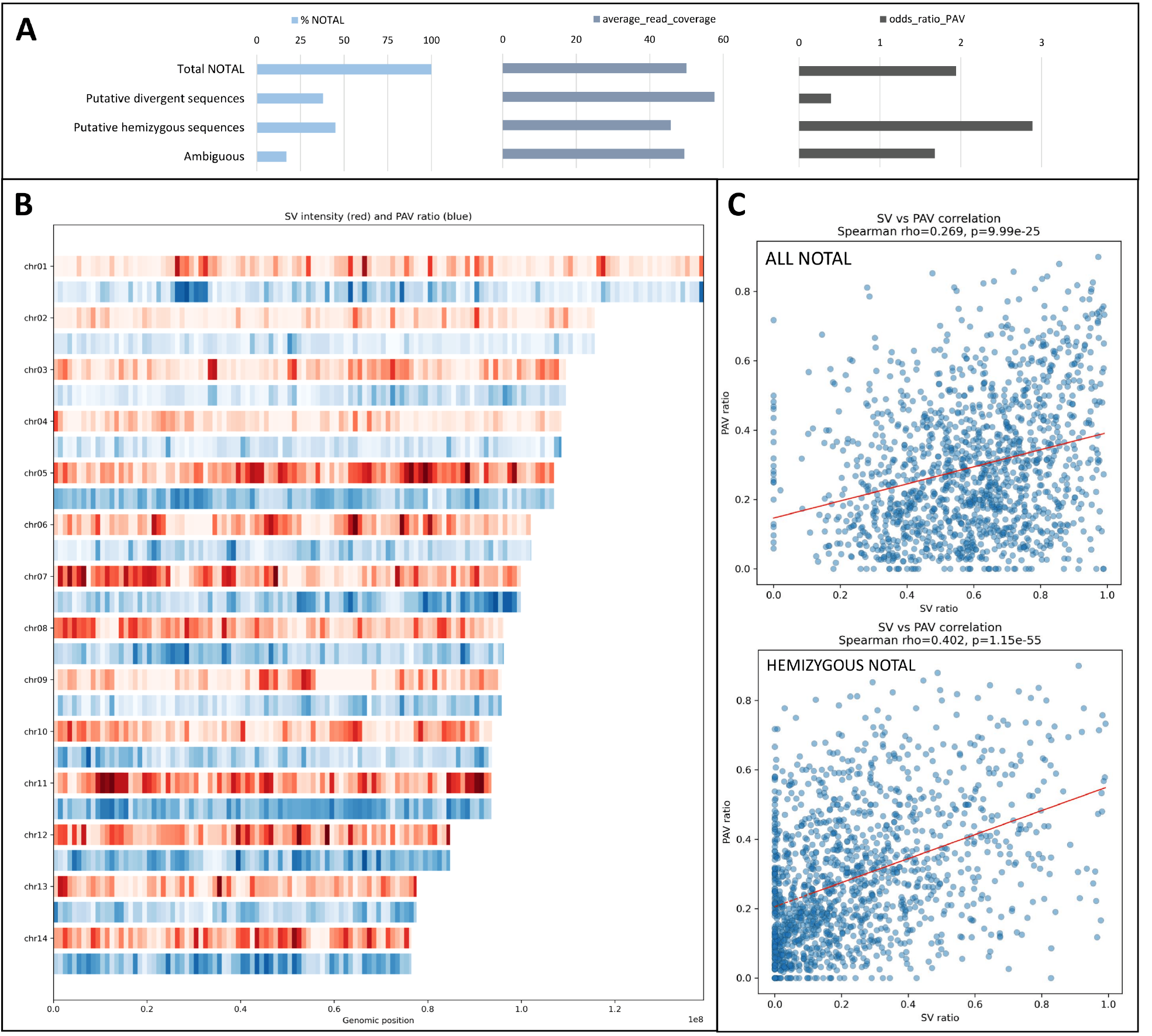
Association between structural variation and dispensable genes. A: Analysis of NOTAL regions distinguishing putative hemizygous sequences (44%) from putative divergent sequences present in both haplotypes (38%). B: Ratio of hemizygous sequences by genomic Mb (red) and ratio of dispensable genes out of total genes per Mb (blue). C: Positive correlations between the locations of dispensable genes and total NOTAL sequences, and the hemizygous filtered fraction, which showed a stronger correlation with dispensable genes than the full NOTAL fraction.

We lastly calculated the distribution of hemizygous sequences and the density of dispensable genes (per Mb of the genome) (Fig. 8B). A significant positive correlation was found between the location of NOTAL regions and dispensable genes, but this was improved when comparing dispensable genes with just the hemizygous fraction filtered out of the total NOTAL fraction (Fig. 8C).

## CONCLUSION

Here we present the first high-quality chromosome-level reference genome for *M. planulatus*, which consists of two fully resolved haplotypes organised into 14 chromosome-scale scaffolds.

Structural comparisons revealed general conservation of synteny between the two *M. planulatus* haplotypes and with *M. edulis* and *M. galloprovincialis*, but also showed that a significant fraction of the genome consists of large hemizygous haplotype-unique NOTAL sequences. These regions are a major component of the pangenomic structure of the *M. planulatus* genome, and genes encoded within them are frequently dispensable. Consistent with this observation, the vast majority of dispensable genes were located in regions associated with SV between haplotypes, as highlighted by the strong positive correlation observed between the density of dispensable genes and hemizygous sequences.

Population structure based on dispensable gene content showed high concordance with SNP-based analyses, that we carried out with a highly conservative approach focused on the polymorphisms associated with core genomic regions shared by all individuals. These analyses confirmed a clear-cut separation between the pure

*M. planulatus* population from Tasmania and the admixed NSW populations, which show evidence of introgression from two distinct Northern *M. galloprovincialis* lineages. Functional enrichment of dispensable genes recovered several gene families associated with transposable element activity, consistent with the role likely played by these mobile elements in shaping the pangenomic architecture in *Mytilus*, as well with immune and stress response, with possible implications for local adaptation.

By providing the first chromosome-scale reference genome for *M. planulatus*, this study fills a major genomic gap within the genus *Mytilus* and lays the foundation for future investigations into the evolution, population genomics and ecological diversification of Southern Hemisphere mussels.

## Data Availability

The data sets supporting the results of this article are available in the NCBI umbrella project under the accession number PRJNA1460848. The raw reads were deposited in the SRA repository under the ac-5 cession numbers SRX32516044 (Hi-C data), SRX32516043 (Pacbio data), and PRJNA1516959 (resequenced individuals). The final genome assembly was deposited in the NCBI Genbank repository, available under the accession numbers: JBXVUX000000000 and JBXVUY000000000 for haplotypes 1 and 2. All additional supporting information, including gene annotations generated by BRAKER for both haplotypes, and the PAV gene matrix, are available in the GigaScience repository, GigaDB [65, 66].

## List of abbreviations

SNP: Single Nucleotide Polymorphism
PAV: presence/absence variation
QV: quality value NOTAL-not-aligned
SV: structural variation
NSW: New South Wales

## Declarations

### Ethics Approval and Consent to participate

The authors declare that ethics approval and consent were not applicable for this research.

## Author Contributions

‐ DK — Conceptualisation, Methodology, Validation, Formal analysis, Investigation, Data curation, Writing — original draft, Writing — review & editing, Visualisation. - DEN — Conceptualisation, Methodology, Validation, Formal analysis, Investigation, Data curation, Writing — original draft, Writing — review & editing, Visualisation. - AS — Conceptualisation, Methodology, Validation, Formal analysis, Investigation, Writing — original draft, Visualisation. - EB — Conceptualisation, Methodology, Validation, Investigation, Writing — review & editing, Visualisation. - SH — Resources, Data curation, Writing — review & editing. - UR — Conceptualisation, Methodology, Writing — review & editing, Supervision. - CR — Resources, Data curation, Writing — review & editing, Supervision. ‐ MG — Conceptualisation, Methodology, Validation, Formal analysis, Investigation, Writing — original draft, Writing — review & editing, Supervision, Project administration, Funding acquisition.

## Appendix

**Table 1a.** Significantly enriched PFAM domains.

| PFAM_ID | Total | Obs. | pvalue | FDR | Bonferroni | Holm-Bonf | Obs-Exp | Description | log10FDR | SL_FDR | direction |
|---|---|---|---|---|---|---|---|---|---|---|---|
| PF00643 | 1071 | 381 | 5.65E-59 | 5.16E-57 | 1.29E-55 | 1.28E-55 | 123.78 | B-box zinc finger | -56.29 | 56.29 | sig_up |
| PF18738 | 531 | 209 | 1.17E-40 | 1.03E-38 | 2.68E-37 | 2.65E-37 | 81.47 | DZIP3/ hRUL138-like HEPN nuclease. Fusion to TPR, Zn-ribbon, RING, Ankyrin, CARD, NACHT ATPase, DEATH and LRR in various animal lineages | -37.99 | 37.99 | sig_up |
| PF00386 | 1149 | 326 | 7.95E-29 | 6.71E-27 | 1.81E-25 | 1.79E-25 | 50.05 | C1q domain | -26.1731 | 26.17309 | sig_up |
| PF12796 | 1384 | 368 | 1.90E-26 | 1.55E-24 | 4.34E-23 | 4.29E-23 | 35.60611 | Ankyrin repeats (3 copies) | -23.81 | 23.80996 | sig_up |
| PF04548 | 207 | 95 | 1.63E-25 | 1.28E-23 | 3.71E-22 | 3.66E-22 | 45.28502 | AIG1 family | -22.8934 | 22.89335 | sig_up |
| PF13359 | 673 | 207 | 1.58E-23 | 1.20E-21 | 3.61E-20 | 3.56E-20 | 45.36627 | DDE superfamily endonuclease | -20.9199 | 20.91989 | sig_up |
| PF00147 | 591 | 188 | 2.14E-23 | 1.57E-21 | 4.88E-20 | 4.81E-20 | 46.06012 | Fibrinogen beta and gamma chains, C-terminal globular domain | -20.8031 | 20.80306 | sig_up |
| PF00619 | 259 | 93 | 3.23E-16 | 2.30E-14 | 7.37E-13 | 7.27E-13 | 30.79623 | Caspase recruitment domain | -13.6378 | 13.63777 | sig_up |
| PF16095 | 248 | 82 | 2.61E-12 | 1.75E-10 | 5.95E-09 | 5.86E-09 | 22.43809 | C-terminal of Roc, COR-A domain | -9.75698 | 9.756977 | sig_up |
| PF05485 | 203 | 70 | 9.36E-12 | 6.10E-10 | 2.13E-08 | 2.10E-08 | 21.24569 | THAP domain | -9.21481 | 9.214806 | sig_up |
| PF00023 | 325 | 97 | 3.06E-11 | 1.94E-09 | 6.98E-08 | 6.87E-08 | 18.94508 | Ankyrin repeat | -8.7125 | 8.712501 | sig_up |
| PF20266 | 320 | 95 | 6.59E-11 | 4.06E-09 | 1.50E-07 | 1.48E-07 | 18.14592 | Mab-21 protein HhH/H2TH-like domain | -8.39105 | 8.391046 | sig_up |
| PF13637 | 473 | 126 | 2.54E-10 | 1.53E-08 | 5.80E-07 | 5.70E-07 | 12.40006 | Ankyrin repeats (many copies) | -7.81646 | 7.816459 | sig_up |
| PF14529 | 641 | 160 | 2.92E-10 | 1.71E-08 | 6.67E-07 | 6.56E-07 | 6.051674 | Endonuclease-reverse transcriptase | -7.76685 | 7.766852 | sig_up |
| PF00653 | 195 | 64 | 6.83E-10 | 3.80E-08 | 1.56E-06 | 1.53E-06 | 17.16705 | Inhibitor of Apoptosis domain | -7.42029 | 7.420288 | sig_up |
| PF12017 | 39 | 22 | 7.77E-10 | 4.22E-08 | 1.77E-06 | 1.74E-06 | 12.63341 | Transposase protein | -7.37478 | 7.374785 | sig_up |
| PF05699 | 183 | 60 | 2.24E-09 | 1.19E-07 | 5.11E-06 | 5.01E-06 | 16.04907 | hAT family dimerisation domain | -6.92537 | 6.925372 | sig_up |
| PF05049 | 38 | 21 | 2.96E-09 | 1.53E-07 | 6.75E-06 | 6.62E-06 | 11.87358 | Interferon-inducible GTPase (IIGP) | -6.81435 | 6.814347 | sig_up |
| PF09588 | 200 | 63 | 5.76E-09 | 2.92E-07 | 1.31E-05 | 1.29E-05 | 14.9662 | YqaJ-like viral recombinase domain | -6.53487 | 6.534867 | sig_up |
| PF13613 | 163 | 54 | 8.20E-09 | 4.07E-07 | 1.87E-05 | 1.83E-05 | 14.85245 | DDE superfamily endonuclease | -6.39074 | 6.390744 | sig_up |
| PF00665 | 181 | 58 | 1.03E-08 | 5.02E-07 | 2.36E-05 | 2.31E-05 | 14.52941 | Integrase core domain | -6.29932 | 6.299323 | sig_up |
| PF05970 | 208 | 64 | 1.25E-08 | 5.95E-07 | 2.85E-05 | 2.80E-05 | 14.04485 | PIF1-like helicase | -6.22574 | 6.225741 | sig_up |
| PF08477 | 195 | 61 | 1.29E-08 | 6.02E-07 | 2.95E-05 | 2.89E-05 | 14.16705 | Miro-like protein | -6.22014 | 6.220137 | sig_up |
| PF12560 | 21 | 14 | 1.65E-08 | 7.52E-07 | 3.76E-05 | 3.68E-05 | 8.956451 | Protein of unknown function (DUF3745) | -6.12357 | 6.123566 | sig_up |
| PF04665 | 355 | 95 | 2.44E-08 | 1.07E-06 | 5.55E-05 | 5.43E-05 | 9.740006 | Poxvirus A32 protein | -5.97134 | 5.971335 | sig_up |
| PF13927 | 434 | 111 | 2.90E-08 | 1.25E-06 | 6.61E-05 | 6.46E-05 | 6.766656 | Immunoglobulin domain | -5.90417 | 5.904174 | sig_up |
| PF07686 | 92 | 35 | 4.27E-08 | 1.80E-06 | 9.74E-05 | 9.51E-05 | 12.90445 | Immunoglobulin V-set domain | -5.74379 | 5.743791 | sig_up |
| PF13873 | 116 | 41 | 4.74E-08 | 1.97E-06 | 0.000108 | 0.000106 | 13.1404 | Myb/SANT-like DNA-binding domain | -5.70659 | 5.706588 | sig_up |
| PF14291 | 86 | 33 | 7.44E-08 | 3.03E-06 | 0.00017 | 0.000166 | 12.34547 | Domain of unknown function (DUF4371) | -5.51832 | 5.518317 | sig_up |
| PF20231 | 166 | 52 | 1.20E-07 | 4.82E-06 | 0.000275 | 0.000268 | 12.13195 | Family of unknown function (DUF6589) | -5.31702 | 5.317021 | sig_up |
| PF13606 | 125 | 42 | 1.78E-07 | 6.99E-06 | 0.000405 | 0.000395 | 11.97888 | Ankyrin repeat | -5.15571 | 5.155708 | sig_up |
| PF03281 | 268 | 73 | 3.77E-07 | 1.46E-05 | 0.000861 | 0.000839 | 8.634709 | Mab-21 protein | -4.83597 | 4.835973 | sig_up |
| PF06739 | 104 | 36 | 4.80E-07 | 1.83E-05 | 0.001095 | 0.001067 | 11.02242 | Beta-propeller repeat | -4.73857 | 4.738567 | sig_up |
| PF00098 | 215 | 61 | 6.41E-07 | 2.40E-05 | 0.001463 | 0.001424 | 9.363666 | Zinc knuckle | -4.62018 | 4.620181 | sig_up |
| PF12248 | 230 | 64 | 7.67E-07 | 2.82E-05 | 0.00175 | 0.001703 | 8.761131 | Farnesoic acid 0-methyl transferase | -4.54928 | 4.549279 | sig_up |
| PF04904 | 29 | 15 | 1.06E-06 | 3.84E-05 | 0.002422 | 0.002356 | 8.035099 | NAB conserved region 1 (NCD1) | -4.41513 | 4.41513 | sig_up |
| PF15135 | 35 | 16 | 4.99E-06 | 0.000172 | 0.011376 | 0.011052 | 7.594085 | Uncharacterised protein UPF0515 | -3.76354 | 3.763545 | sig_up |
| PF20209 | 150 | 44 | 6.66E-06 | 0.000227 | 0.015202 | 0.014763 | 7.974651 | Domain of unknown function (DUF6570) | -3.64416 | 3.644162 | sig_up |
| PF02945 | 87 | 29 | 1.14E-05 | 0.000383 | 0.026035 | 0.025271 | 8.105297 | Recombination endonuclease VII | -3.41695 | 3.416947 | sig_up |
| PF13920 | 243 | 63 | 1.18E-05 | 0.00039 | 0.026892 | 0.02609 | 4.638934 | Zinc finger, C3HC4 type (RING finger) | -3.40923 | 3.40923 | sig_up |
| PF06140 | 37 | 16 | 1.28E-05 | 0.000416 | 0.029116 | 0.028235 | 7.113747 | Interferon-induced 6-16 family | -3.38097 | 3.380967 | sig_up |
| PF00078 | 2741 | 562 | 1.73E-12 | 1.20E-10 | 3.95E-09 | 3.89E-09 | -96.3032 | Reverse transcriptase (RNA-dependent DNA polymerase) | -9.9222 | -9.9222 | sig_down |
| PF13855 | 876 | 206 | 3.23E-10 | 1.84E-08 | 7.37E-07 | 7.24E-07 | -4.38804 | Leucine rich repeat | -7.73477 | -7.73477 | sig_down |
| PF00059 | 1206 | 259 | 2.34E-08 | 1.05E-06 | 5.34E-05 | 5.22E-05 | -30.6438 | Lectin C-type domain | -5.98035 | -5.98035 | sig_down |
| PF01582 | 563 | 130 | 1.20E-06 | 4.29E-05 | 0.002747 | 0.002671 | -5.21514 | TIR domain | -4.36733 | -4.36733 | sig_down |
| PF00096 | 1017 | 214 | 1.54E-06 | 5.40E-05 | 0.00351 | 0.003412 | -30.2519 | Zinc finger, C2H2 type | -4.26755 | -4.26755 | sig_down |

**Figure 1a.**
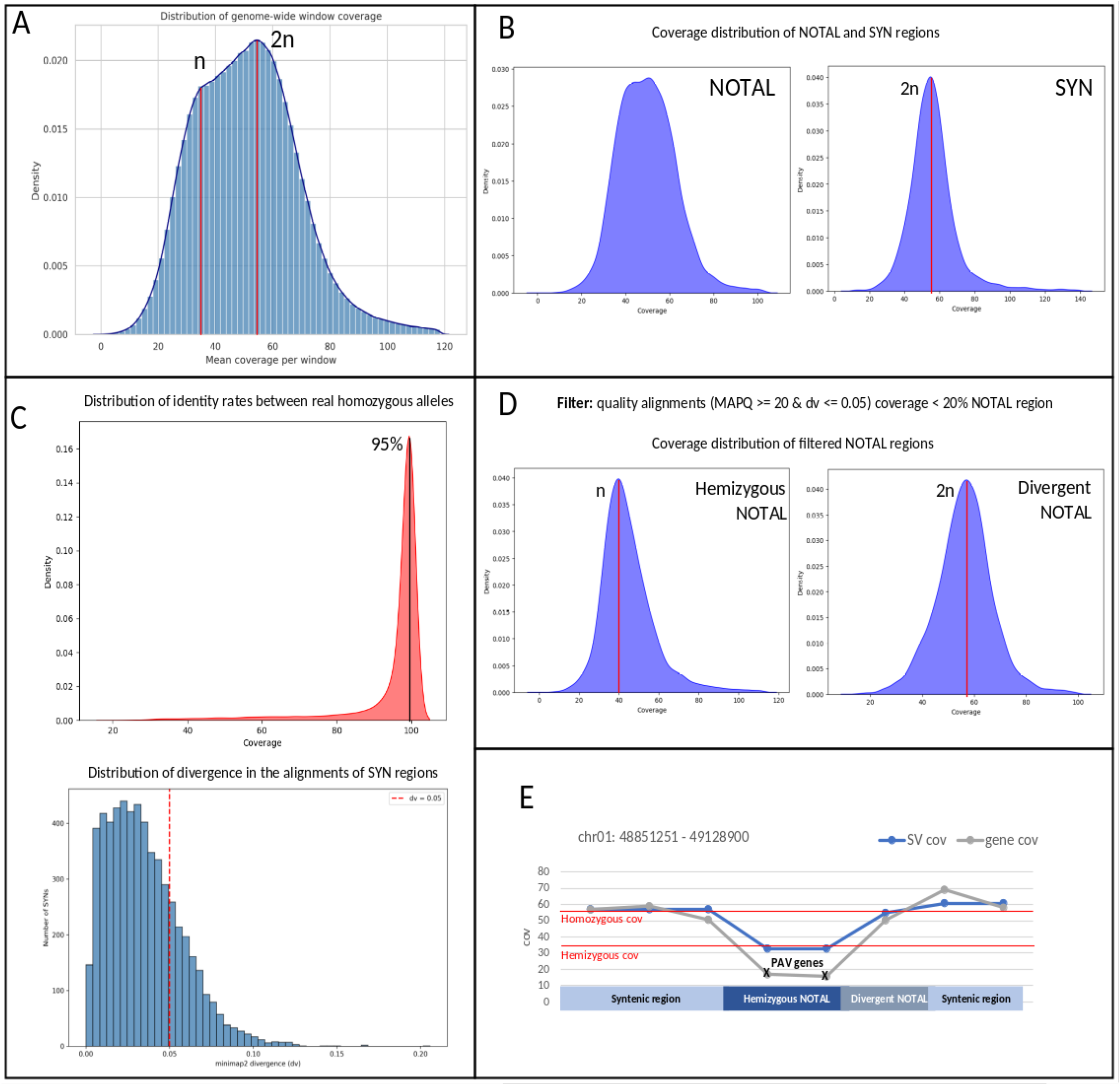
Analysis of structural variation between haplotypes. A. Coverage distribution in 1 kb windows across the Mytilus planulatus genome. The homozygosity signal is placed at the expected coverage (2n), while we see another hemizygous signal which would have approximately half of the expected coverage (n). This kind of patterns are typical in bivalve genomics. B. The coverage distribution of syntenic regions (SYN) across haplotypes is homozygous, as expected. The coverage distribution of the not aligned regions between haplotypes (NOTAL) is clearly not only hemizygous, so these regions will be filtered to separate the real hemizygous fraction. C. The real alleles in the syntenic regions of both haplotypes show an average homology of 95%. That same threshold (0.05 divergence) is shown in the divergence distribution of SYN alignments between haplotypes. The 0.05 divergence threshold is established as the parameter to consider an alignment of sufficient quality, since it is associated with real alleles in synteny analyses. D. NOTAL regions are filtered and the ones showing less than 20% coverage by good quality alignments are considered hemizygous. The coverage distribution supported our filtering strategy, since now the hemizygous NOTAL regions show hemizygous coverage. E. An example selected from chromosome 1 shows syntenic regions with homozygous genes and expected coverage, interrupted by a structural variant consisting in an hemizygous regions with half expected coverage and dispensable genes, entirely absent in re-sequencing data.

